## Supplementary File 1 for "An engineered biosensor enables dynamic aspartate measurements in living cells"

Supplementary information: Protein engineering and in vitro characterization.

**iAspSnFR2, annotated sequence**

Mutations conferring aspartate specificity (S27A and S72P) was inserted into a precursor of iGluSnFR3 (iGluSnFR3-pre) to obtain iAspSnFR2.

Numbering according to 2VHA.pdb (glutamate binding protein, purple) and 2B3P.pdb (superfolder GFP, orange). Differences between iGluSnFR3.v857, iGluSnFR3-pre and iAspSnFR2 are:

GltI: E25K in iGluSnFR3-pre/iAspSnFR2, E25D in iGluSnFR3.v857

GltII: S27A in iAspSnFR2, S27S in iGluSnFR3-pre/iGlu3SnFR3.v857

GltI: S72P in iAspSnFR2, S72S in iGluSnFR3-pre/iGluSnFR3.v857

GltI: A184V in iGluSnFR3-pre/iAspSnFR2, A184S in iGluSnFR3.v857

cpSFVenus: S147S in iGluSnFR3-pre/iAspSnFR2, S147N in iGluSnFR3.v857

cpSFVenus: N147I in iGluSnFR3-pre/iAspSnFR2, N147T in iGluSnFR3.v857

cpSFVenus: T228 in iGluSnFR3-pre/iAspSnFR2, G228 in iGluSnFR3.v857

cpSFVenus: L46 in iGluSnFR3-pre/iAspSnFR2, F46 in iGluSnFR3.v857

cpSFVenus: GYG chromophore in iGluSnFR3-pre/iAspSnFR2, SYG chromophore in iGluSnFR3.v857

```
         1         2         3         4         5         6
         0         0         0         0         0         0
 MRSAAGSTLDKIAKNGVIVVGHRKSAVPFSYYDSQHKVVGYSQEYSNAIVEAVKKKLNK

                                       1         1         1
         7         8         9         0         1         2
         0         0         0         0         0         0
PDLQVKLIPITPQNRIPLLQNGTYDFECGSTTNNVERQKQAAFSDTIFVVGTRLLTKKGG


         1         1         1         1         1         1
         3         4         5         6         7         8
         0         0         0         0         0         0
DIKDFANLKDKAVVVTSGTTSEVLLNKLNEEQKMNMRIISAKDHGDSFRTLESGRAVAFM

         1         2         2         2         2         2
         9         0         1         2         3         4
         0         0         0         0         0         0
MDDVLLAGERAKAKKPDDWEIVGKPQSQEAYGCMLRKDDPQFKKLMDDTIAQVQTSGEAE

         2   1     1         1         1         1         1
         5   4     5         6         7         8         9
         0   7     3         3         3         3         3
KWFDKWFKNPILVSHIVYITADKQKNGIKANFKIHHNVEDGSVQLADHYQQNTPIGDGPV

         2         2         2         2    2
         0         1         2         3    3
         3         3         3         3    8      1       9
LLPDNHYLSYQSVLSKDPNEKRDHMVLLEFVTATTNSLGMDELYKGGTGGSMSKGEELFT


         1         2         3         4         5     666
         9         9         9         9         9     567
GVVPILVELDGDVNGHKFSVRGEGEGDATNGKLTLKLICTTGKLPVPWPTLVTTLGYGVQ

                                       1         1         1
         7         8         9         0         1         2
         9         9         9         9         9         9
CFARYPDHMKQHDFFKSAMPEGYVQERTISFKDDGTYKTRAEVKLEGDTLVNRIELKGID


         1      1      2         2         2
         3      4      6         7         7
         9      6      0         0         9
FKEDGNILGHKLEYNFNNPLNMNFELSDEMKALFKEPNDKALKLQHHHHHH
```

**Pairwise sequence alignment of iAspSnFR (Hellweg et al., 2023) and iAspSnFR2 (this work)**

Pairwise sequence alignment using the GFP and GltI domains only. GltI domain in purple, GFP in orange. Mutations conferring aspartate specificity to the iGluSnFR3 precursor (S27A and S72P) marked by yellow background.

Difference between iAspSnFR and iAspSnFR2 indicated by colon. A total of 25 mismatches and 497/522 = 95.2% sequence identity.

```
iAspSnFR     1 ---AAGSTLDKIAKNGVIVVGHRESSVPFSYYDNQQKVVGFSQDYSNAIV  47
                  ||||||||||||||||||||:|:|||||||:|:||||:||:||||||
iAspSnFR2    1 MRSAAGSTLDKIAKNGVIVVGHRKSAVPFSYYDSQHKVVGYSQEYSNAIV  50

iAspSnFR    48 EAVKKKLNKPDLQVKLIPITAQNRIPLLQNGTFDFECGSTDNNVERQKQA  97
               ||||||||||||||||||||:|||||||||||:|||||||:|||||||||
iAspSnFR2   51 EAVKKKLNKPDLQVKLIPITPQNRIPLLQNGTYDFECGSTTNNVERQKQA 100

iAspSnFR    98 AFSDTIFVVTTRLLTKKGGDIKDFANLKDKAVVVTSGTTSEVLLNKLNEE 147
               |||||||||:||||||||||||||||||||||||||||||||||||||||
iAspSnFR2  101 AFSDTIFVVGTRLLTKKGGDIKDFANLKDKAVVVTSGTTSEVLLNKLNEE 150

iAspSnFR   148 QKMNMRIISAKDHGDSFRTLESGRAVAFMMDDVLLAGERARAKKPDNWEI 197
               ||||||||||||||||||||||||||||||||||||||||:|||||:|||
iAspSnFR2  151 QKMNMRIISAKDHGDSFRTLESGRAVAFMMDDVLLAGERAKAKKPDDWEI 200

iAspSnFR   198 VGKPQSQEAWGCMLRKDDPQFKKLMDDTIAQVRTSGEAEKWFDKWFKNPI 247
               |||||||||:||||||||||||||||||||||:|||||||||||||||||
iAspSnFR2  201 VGKPQSQEAYGCMLRKDDPQFKKLMDDTIAQVQTSGEAEKWFDKWFKNPI 250

iAspSnFR   248 LVSHNVYITADKQKNGIKANFKIRHNVEDGSVQLADHYQQNTPIGDGPVL 297
               ||||:||||||||||||||||||:||||||||||||||||||||||||||
iAspSnFR2  251 LVSHIVYITADKQKNGIKANFKIHHNVEDGSVQLADHYQQNTPIGDGPVL 300

iAspSnFR   298 LPDNHYLSTQSVLSKDPNEKRDHMVLLEFVTAAGITLGMDELYKGGTGGS 347
               ||||||||:|||||||||||||||||||||||::::||||||||||||||
iAspSnFR2  301 LPDNHYLSYQSVLSKDPNEKRDHMVLLEFVTATTNSLGMDELYKGGTGGS 350

iAspSnFR   348 MSKGEELFTGVVPILVELDGDVNGHKFSVRGEGEGDATNGKLTLKFICTT 397
               |||||||||||||||||||||||||||||||||||||||||||||:||||
iAspSnFR2  351 MSKGEELFTGVVPILVELDGDVNGHKFSVRGEGEGDATNGKLTLKLICTT 400

iAspSnFR   398 GKLPVPWPTLVTTLTYGVQCFSRYPDHMKQHDFFKSAMPEGYVQERTISF 447
               ||||||||||||||:||||||:||||||||||||||||||||||||||||
iAspSnFR2  401 GKLPVPWPTLVTTLGYGVQCFARYPDHMKQHDFFKSAMPEGYVQERTISF 450

iAspSnFR   448 KDDGTYKTRAEVKFEGDTLVNRIELKGIDFKEDGNILGHKLEYNFNNPLN 497
               |||||||||||||:||||||||||||||||||||||||||||||||||||
iAspSnFR2  451 KDDGTYKTRAEVKLEGDTLVNRIELKGIDFKEDGNILGHKLEYNFNNPLN 500

iAspSnFR   498 MNFELSDEMKALFKEPNDKALK 519
               ||||||||||||||||||||||
iAspSnFR2  501 MNFELSDEMKALFKEPNDKALK 522
```
